# Neural entrainment to pitch changes of auditory targets in noise

**DOI:** 10.1101/2025.04.03.646585

**Authors:** Xiaoxuan Guo, Guangting Mai, Yousef Mohammadi, Ester Benzaquén, Kate Yukhnovich, Will Sedley, Timothy D Griffiths

## Abstract

Neural entrainment to certain acoustic features can predict speech-in-noise perception, but these features are difficult to separate. We measured neural responses to both natural speech-in-noise and stimuli (auditory figure-ground) that simulate speech-in-noise without any acoustic or linguistic confound such as stress contour and semantics. The figure-ground stimulus is formed by multiple temporally coherent pure-tone components embedded in a random tone cloud. Previous work has shown that discrimination of dynamic figure-ground based on the fundamental frequency (F0) of natural speech predicts speech-in-noise recognition independent of hearing and age. In this study, we compared the brain substrate for the figure-ground analysis based on the F0 contour and a statistically similar ‘1/f’ contour with speech-in-noise. We used the temporal response function to predict the electroencephalography responses to the frequency trajectories of the auditory targets. We demonstrate that the brain significantly tracked the pitch changes in both AFG conditions (F0 and 1/F tracking) and a sentence-in-noise condition (F0 tracking) at similar latencies, but at similar magnitudes only when tracking the F0 contour. The pitch-tracking accuracy was consistently high across the delta and theta bands for the AFG condition but not for speech. Sensor-space analysis revealed that speech-in-noise performance correlated with the positive peak amplitude of the F0 figure-ground at 100 ms. Source-space analysis revealed bilateral temporal lobe and hippocampal generators, and strong tracking in the superior parietal lobe for auditory figures and natural speech. In conclusion, our findings demonstrate that the human brain reliably tracks the F0 trajectory of both speech and a non-linguistic figure in noise, with speech tracking showing reduced accuracy in the theta band compared to figure-ground tracking. Despite the difference in prediction accuracy, we reveal striking similarities in neural entrainment patterns and source locations between the two paradigms. These results suggest that neural entrainment engages high-level cortical mechanisms independent of linguistic content. Furthermore, we show that TRF peak amplitude serves as a potential biomarker for speech-in-noise ability, highlighting possible clinical applications.

## Introduction

Segregating and tracking a target sound in complex acoustic environments is an important skill that the auditory system performs to facilitate daily activities (Cherry, 1953; McDermott, 2009). When segregating speech from a noisy environment, humans rely on auditory and cognitive mechanisms to process target speech that stands out due to its acoustic features, even before any language processing (Lad et al., 2024; Guo et al., 2022; Finkl et al., 2020; Oster & Werner, 2020). These features include frequency and temporal cues, source location and timbre. Segregation is continuous in the natural environment and engages auditory cognitive mechanisms including perception, working memory and attention (Akeroyd, 2008; Shinn-Cunningham & Best, 2008). In this work we seek to elucidate neural correlates of segregation using stimuli with similar complexity to speech but in the absence of high-level linguistic information. This allows a comparison between pre-linguistic mechanisms for segregation and speech-in-noise (SIN) perception. The work has the potential to suggest a language-independent measure to explain SIN deficits that are not accounted for by peripheral deafness.

The current study depends on the further development of an Auditory Figure-Ground (AFG) task based on a stimulus with fixed-frequency components that assesses auditory segregation relevant to SIN perception (Teki et al., 2011, 2013). The prototype AFG is called the stochastic figure-ground, comprising an auditory figure made of fixed-frequency temporally invariant tones, and a ground comprising random tones. Modelling work suggests a figure-tracking mechanism based on the detection of temporal coherence between the component frequencies (Teki et al 2016), which was also evidenced by an electrophysiology work (O’Sullivan et al., 2015a) that demonstrated neural tracking of the coherence level of the auditory figure. Brain imaging studies support cortical mechanisms beyond the primary cortex that overlap with those for SIN (Guo et al., 2022; Holmes et al., 2021; Holmes & Griffiths, 2019; O’Sullivan et al., 2015; Schneider et al., 2018; Teki et al., 2016a, 2016b).

While fixed-frequency figure-ground was shown to measure the fundamental sound grouping aspect of SIN processing, natural speech has richer information embedded. One of the most perceptually salient features of natural speech is the pitch, the value of which is determined by the fundamental frequency of voiced sounds. Pitch perception plays a crucial role in sound segregation (D’Alessandro et al., 2024; Oxenham, 2008), and training in pitch discrimination improves SIN performance (Gohari et al., 2023; Moossavi et al., 2021). However, pitch contour is highly correlated with other aspects of speech prosody (rhythm and stress contour), making it difficult to isolate the effect of pitch processing in a natural auditory scene containing speech. To address this issue, we have developed a dynamic auditory figure-ground paradigm that simulates the pitch changes in natural sentences based on a stimulus with isolated pitch changes and no linguistic confounds (Guo et al., 2024). The aim was to measure the behavioural performance and brain substrates for a ‘pure’ type of pitch tracking in noise as an important component of SIN perception.

The dynamic AFG stimulus assesses brain mechanisms for tracking a pitch contour derived from speech. Our previous behavioural work showed that dynamic AFG based on the trajectory of fundamental frequency (F0) in human speech (AFG-F0) predicted a large variance of the SIN performance at both the word and sentence level in a multivariate model including hearing sensitivity, age, and both static and dynamic figure-ground tasks (Guo et al., 2024). However, we do not yet know if the brain parses the AFG information the same way as SIN and if it can reliably track F0 in the AFG stimulus as in natural speech. In this study, we investigate the neural entrainment to both SIN and AFG-dynamic by analysing the EEG temporal response function (TRF) of the frequency profiles embedded in the stimuli.

Neural entrainment here refers to the temporal alignment of neural oscillations with acoustic regularities of a sound. TRF is a method that has been used to investigate the linear mapping of EEG signals to sound features (Obleser & Kayser, 2019). TRF captures more precise characterization of sensory responses to naturalistic speech stimuli than simple correlations between neural and speech signals as cross-correlation can cause temporal smearing when analysing slow-changing stimuli, such as speech, which correspond to the EEG signal at multiple overlapping time lags (Crosse et al., 2016). To further dissect if the entrainment can only be evoked by natural speech pitch patterns or any speech-like frequency contours, we also included a condition with AFG following the 1/F trajectory (AFG-1/F). 1/F fluctuation has been observed in speech and music, and it was hypothesised to reflect a fundamental aspect of physiological and cognitive functions (Kello et al., 2007; Voss & Clarke, 1975). It is a speech-like signal without linguistic prosody or the semi-periodic frequency contour of sentences, which provides further control of the linguistic information in the AFG stimuli. In addition to EEG sensor-level analysis, the source locations of the TRF peak responses are investigated in the current work to study if the neural generators of the dynamic AFG are similar to that of SIN compared to the previous neuroimaging studies of static AFG and SIN (Holmes et al., 2021; Teki et al., 2016). We hypothesised that the brain would track the frequency changes in the pitch contour of speech and figure similarly, in terms of the TRF spectrotemporal features and source locations when masked by noise.

This paradigm has potential clinical applications. Currently, available behavioural SIN tests, such as QuickSIN, SCAN-3C, or LiSN-S, rely heavily on verbal responses (Browne et al., 2024; Cameron & Dillon, 2007). These tests therefore exclude people who are not able to give reliable responses (e.g., due to language production deficits or developmental disorders and cognitive impairments). EEG recordings of SIN responses (Panela et al., 2024; Guo et al., 2022; Muncke et al., 2022) allow a measure of brain activity that is not dependent on response. Here, we seek EEG responses to a more fundamental level of auditory processing before linguistic analysis. The work has the potential to isolate ‘intermediate’ mechanisms for SIN relevant to speakers of any language with any degree of proficiency, between the level of cochlear processing (measured with the audiogram or otoacoustic emissions) and actual speech in noise (measured with speech stimuli).

## Methods

### Participants

Thirty-four participants attended the recording sessions, of whom one was excluded due to poor EEG recording quality, and one non-native English speaker was excluded. The sample size was deemed sufficient based on a previous study with a similar design, which included 10 participants (O’Sullivan et al., 2015b). The full inclusion criteria were as follows: participants should be native English speakers with no history of auditory, language, psychological, developmental or neurological disorders, and who are not currently taking any psychotropic drugs. People with mild hearing loss were included as long as they were able to perform all the tests. Normal hearing is defined as the pure-tone average under 25 dB across 250 Hz, 500 Hz, 1000 Hz, 2000 Hz, and 4000 Hz, and mild hearing loss is defined as 26 – 40 dB (Olusanya et al., 2019). Participants between 18 and 70 were included for a more representative sample. The final analysis was carried out on 32 participants (13 women) aged from 22 to 67 (mean = 40.19, standard deviation (SD) = 13.68). The pure-tone audiogram (PTA) results are shown in Figure 1. This study followed the Helsinki ethical standards and was approved by the Newcastle University Ethics Committee (46225/2023).

**Figure 1.**
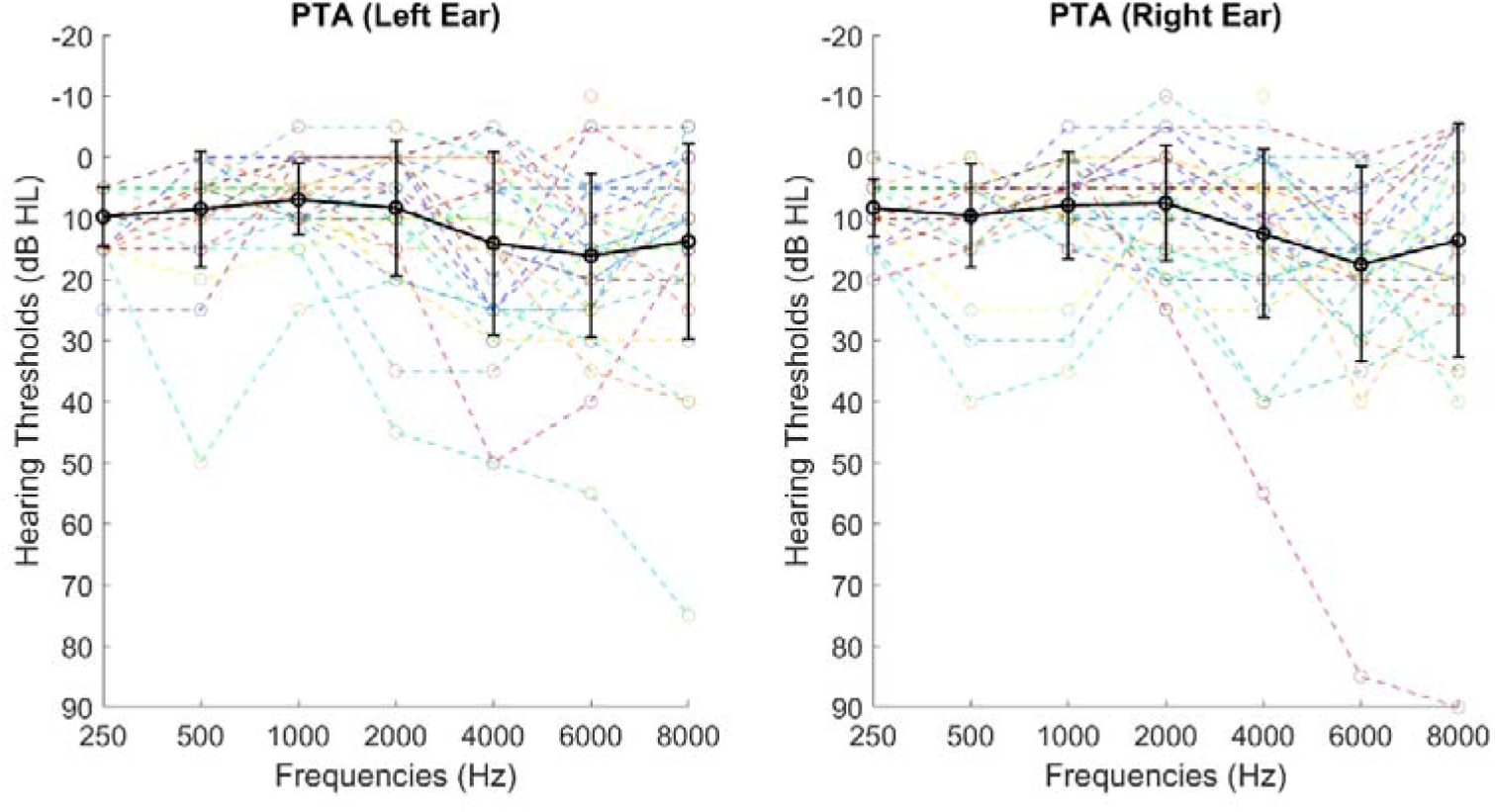
Average pure-tone audiogram results of 250 - 8000 Hz of all participants in dashed lines of multiple colours. The thick black line plots the group average with standard deviation bars.

### Stimuli and Experimental Design

The AFG-F0 stimuli were adapted from Guo et al. (2024). The auditory target follows the trajectory of the fundamental frequency of speech sentences (Figure 2a) taken from the SIN task from Holmes & Griffiths (2019) with a frequency range of 74.94 - 295.44 Hz (M=131.59, SD=15.61) using Praat (Boersma, 2001). Any gaps in the frequency contours were removed. The signals were then detrended and lowpass filtered at 3 kHz to remove sharp transitions that would otherwise be a strong cue for perception (Figure 2a). The frequency elements were 50 ms each and they were concatenated to form a continuous trajectory. The fundamental trajectory was multiplied by 2, 3, and 4 to form a harmonic structure (see Figure 2b). These figures were masked by a tone cloud of 10 elements per time point with pseudo-randomly generated frequency elements in the ground from around 90 Hz to around 3623 Hz following a logarithmic scale. The ground stimuli were constrained to have no overlapping frequency elements with the figure. The figure and the ground have the same onset and offset and were played at the same sound intensity level across participants (target-to-masker ratio, or TMR at 0 dB). This ensured that the segregation of the figure from the ground relied strictly on the temporal coherence of the figure as defined by Teki et al. (2016). Each segment of the figure-ground was then concatenated sequentially to make a longer continuous sound that lasted around 9 minutes per trial.

**Figure 2.**
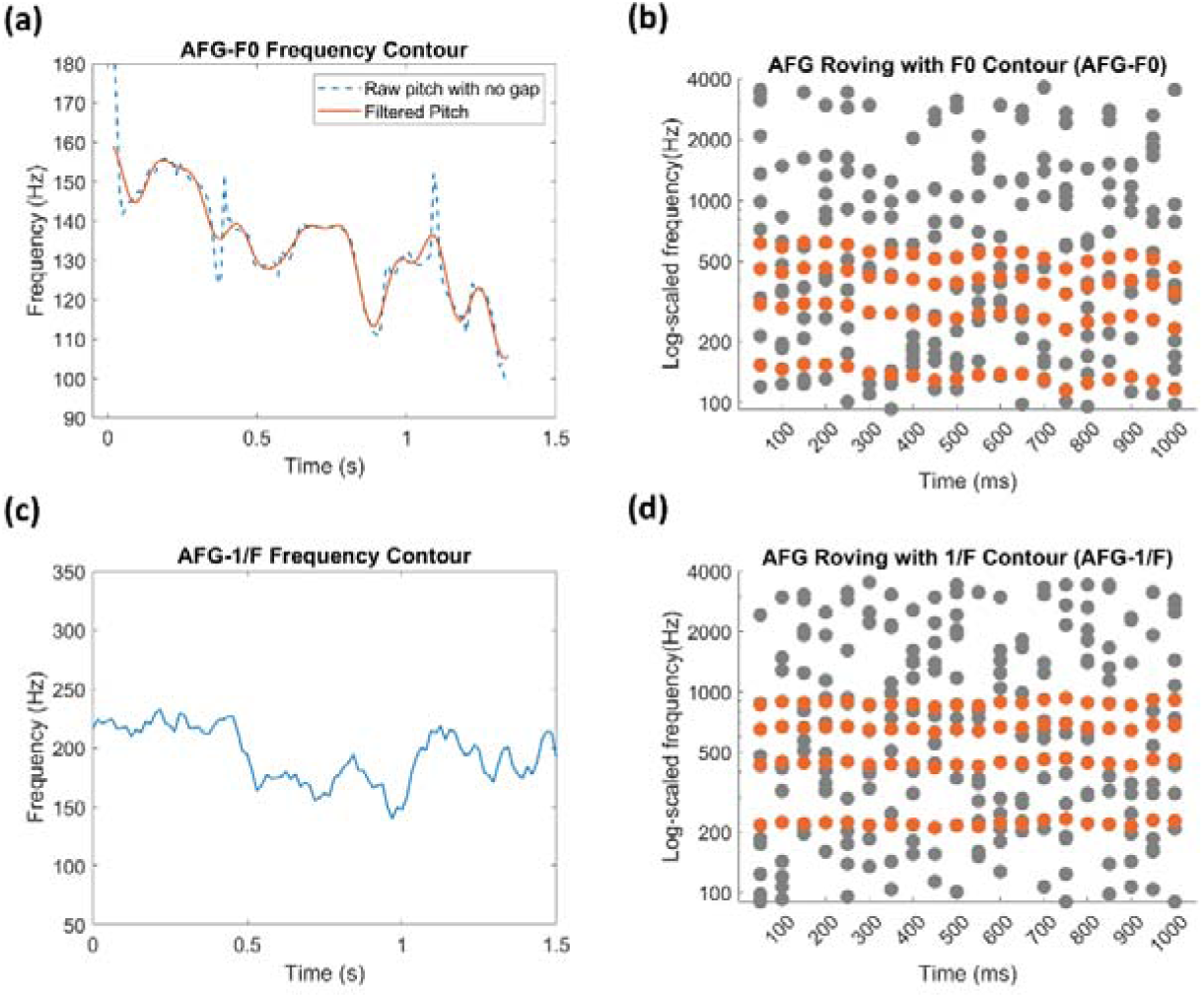
The frequency contours of AFG-F0 (Figure 2(a)) and 1/F (Figure 2(c)) and the figure-ground dotted plots (Figure 2(c)(d)). The x-axes of Figure 2(a)(c) are time durations in seconds. The y-axes are the frequencies in Hz. Figure 2 (a) shows the raw pitch (the same as in actual speech) in dotted lines and the filtered pitch contour in red line. Figure 2(b)(d) shows examples of AFG-F0 and AFG-1/F respectively. The red dots plot the auditory figure elements, and the grey dots plot the ground elements. The x-axes in Figure 2 (b)(d) are time durations in milliseconds. The y-axes represent frequencies in Hz.

The figure and ground elements of AFG-1/F stimuli were generated in the same way as the AFG-F0 condition but following artificial pitch trajectories. Briefly, the contour of the 1/F conditions was generated in the frequency domain using a 1/F power spectrum and random phase spectrum. Inverse Fourier transform was performed to obtain the 1/f noise trajectory in the time domain. The frequency series was then normalised and scaled to 74 - 295 Hz to be close to the human speech range. Figure 2c and 2d illustrate an example of the 1/f contour and the AFG-1/F stimulus.

The trial structure is illustrated in Figure 3. The AFG tests consisted of two identical runs presented sequentially with 2 blocks in each trial separated by a self-paced break. The participants were also given a self-paced break between the two trials. Within each block, there were gaps randomly placed in the continuous figure, whilst the ground stimuli continued uninterrupted. These gaps lasted for 600 ms and were randomly placed throughout the testing with each trial consisting of 30 gaps. Participants were asked to identify the gaps and press a button when they could detect a gap. They would need to be able to segregate the figure from the ground continuously during the experiment in order to perform the task, as there was no gap in the ground. This active task was designed to keep the participants’ attention level high throughout the recording to maximise the EEG responses. The length of the stimuli across conditions was kept at a similar length (difference in seconds to keep the integrity of the sentence). Examples of these sounds can be accessed online (https://osf.io/c4rns/files/osfstorage#).

**Figure 3.**
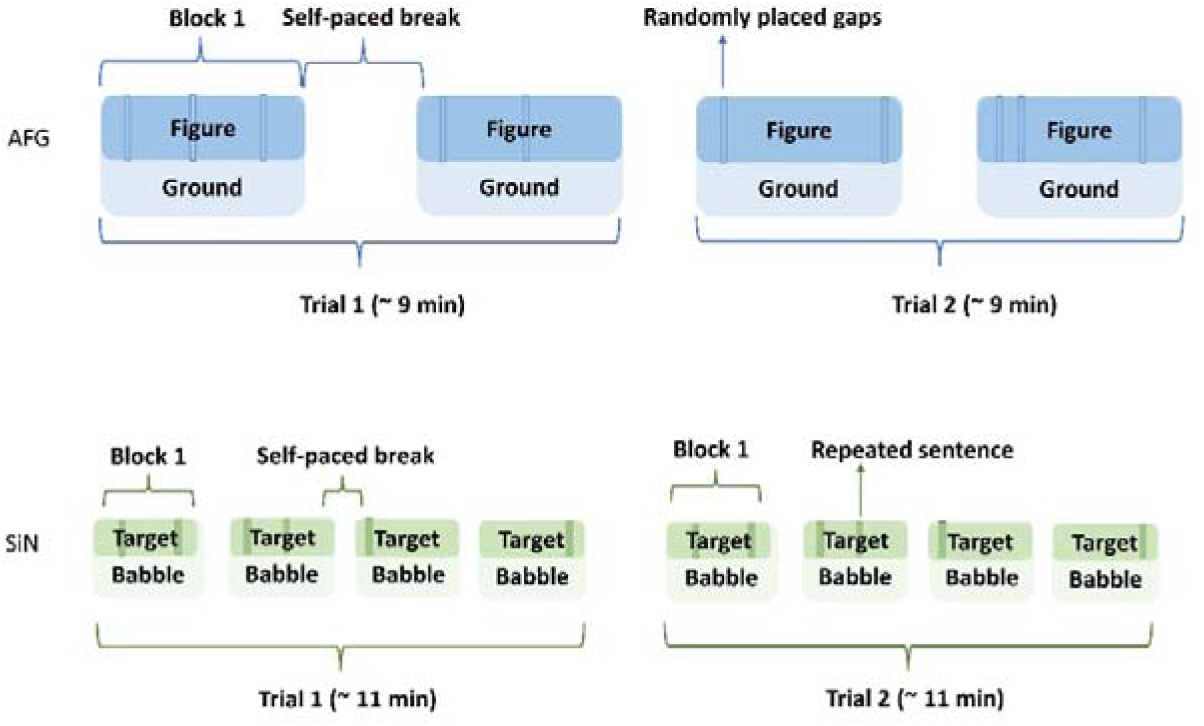
Schematics of experiment design. The top plots show the trial structure of the two AFG conditions. The darker blue rectangles are the figures, and the lighter blue are the grounds. The bottom plots show the trial structures for the SIN condition with the darker green as the target sentences and the lighter green the babble noises.

The SIN stimuli were English versions of the Oldenburg sentences (Holmes & Griffiths, 2019). The target sentences consisted of five words following a structure of Name-Verb-Number-Adjective-Noun (e.g., “Alan ordered twelve white sofas”), which were masked by a 16-talker babble. The signal-to-noise ratio was 0 dB. The sentences had the same onset and offset as the background babble to make SIN segments, which were joined together similarly to the AFG stimuli to form a continuous sound. This was done to simulate a naturalistic conversation flow without giving too many semantic and pragmatic cues to the participants.

The SIN condition contained two identical runs, each containing 4 blocks separated by 3 self-paced breaks. There was also an active task for the SIN condition, in which 30 repeated sentences were randomly placed as the response trials. Participants were asked to press the button when they could detect a repeated sentence. The different active tasks for SIN and AFG conditions were designed based on feasibility. Speech contains natural gaps, making the gap-detection task unfeasible. Similarly, a repeated pattern in the continuous AFG stimuli was too difficult to detect, rendering a pattern discrimination design unfeasible. The number of response trials was thus limited to 30 to minimise behavioural effects on the two conditions while maintaining participants’ attention.

All stimuli were generated off-line with MATLAB R2021a and presented with Psychtoolbox version 3.0.19 through headphones (Sennheiser HD 380 Pro) linked to a sound card (RME FireFace UC).

### Procedure

After giving informed consent, participants were taken to a sound-proof booth for an audiometric test. They were then prepared for the EEG recording session. The researcher briefly explained the tasks and specifically asked the participants to pay attention to the target sound throughout the recording. The task instructions were shown on an LCD display. A fixation cross was displayed at the centre of the screen during all three tasks, and participants were told to fixate their gaze on the cross. Feedback was provided whenever participants pressed a button (both for false alarm and correct detection). The EEG session had three tasks following the same order of presentation across participants: the AFG-F0 gap detection task, SIN repetition detection task, and the AFG-1/F gap detection task. Each task took around 18-25 minutes depending on the duration of the self-paced breaks. Before each experiment, participants were given some example sounds to familiarise themselves with the test stimuli as well as some practice trials that were different from the main experiment. The practice was repeated if the participants failed to do the task until they showed good performance.

Participants were also given longer breaks between tasks with refreshments to minimise fatigue.

### Data Analysis

#### 2.4.1 Psychophysics

D prime (d’) was used to quantify the performance, which was calculated as: d’ = z(hit) – z (false alarm). The extreme values (0% hit rate or false alarm rate) were replaced by 0.5/n or rates of 100% with (n−0.5)/n where n is the number of signal or noise trials (Macmillan & Kaplan, 1985).

#### 2.4.2 Extracting and Processing the Pitch Information

The input pitch information was processed differently in different conditions. For the two AFG conditions, the frequency information was retained in full, including the gaps. The number of gaps was relatively low and should not introduce significant distortion to the results. The 30 gaps were filled by the frequency value before the gap onset. We then took the absolute values of the first derivatives of the F0 or 1/F contours to quantify the absolute pitch change of the auditory stimuli (not taking the absolute value would make the assumption that neural responses to pitch decreases were equal and opposite to pitch increases). These were then resampled and aligned to the EEG data.

For the SIN condition, the extraction of F0 contours followed the same method used to create the AFG-F0 figure. As the speech condition also contains the stress contour (specifically the amplitude envelope), which can confound the EEG responses to pitch, it was regressed out from the raw pitch. We took the residual of the linear regression of F0 on the absolute value of the stress contour extracted through Hilbert transform. This was performed on the target speech only.

Finally, the absolute value of the first derivatives of the processed pitch values of all three conditions was taken as the final stimuli aligning to the EEG signals.

#### 2.4.3 EEG Preprocessing

EEG data acquisition was carried out with a 64-channel BioSemi ActiveTwo system. Data analysis was conducted using MATLAB R2021a with EEGLAB version 2019. The continuous EEG data were first referenced to P9 and P10 (positioned similarly to the mastoids). They were then highpass filtered at 0.1 Hz with a 3^rd^ order Butterworth filter, notch filtered at 49-51 Hz to remove the power line noise, and then lowpass filtered at 30 Hz with the same filter. Following the filtering, intervals where participants took long breaks were removed from the data first to increase the processing speed. The remaining data were downsampled from 2048 Hz to 100 Hz. The Artifact subspace reconstruction (ASR) tool was used to detect noisy channels: channels poorly correlated (r<0.6) with their random sample consensus reconstruction were rejected and interpolated. Independent component analysis (ICA) was applied to remove artifacts such as eye blinks, muscle activity, and heart rate. Up to 16 components were excluded based on visual inspection and classification using the EEGLAB IClabel extension (Pion-Tonachini et al., 2019). After ICA component rejection, the EEG data were re-referenced to a common average reference, following which the data were epoched into 4 epochs for the AFG conditions and 8 epochs for the SIN condition corresponding to their respective experimental blocks. The Cz channel was chosen for analysis based on a previous study, where the researchers found that vertex-to-mastoid analysis could show reliable responses of figure-ground segregation (Guo et al., 2022).

Finally, EEG data were filtered to delta (1-4 Hz) and theta (4-8 Hz) frequency bands using a Butterworth bandpass filter (3^rd^ order). These frequency ranges were chosen as the current study investigated the pitch changes, which is a low-changing frequency contour. The delta and theta frequency bands are relevant for tracking pitch-related information and have often been used to study the low (prosody) and higher-level (syntactic) features of speech (Mai & Wang, 2023; Etard & Reichenbach, 2019; Behroozmand et al., 2015; Giraud & Poeppel, 2012). Furthermore, the AFG stimuli contain pure-tone elements that occur regularly every 50 ms, which would elicit 20 Hz neural entrainment and confound the analysis for the fundamental frequency for a higher frequency range.

#### 2.4.4 Computing Temporal Response Forward Model

The temporal response function (TRF, Kegler et al., 2022; Lalor et al., 2006) was used to analyse the relationship between pitch and the EEG responses using the mTRF-Toolbox (Crosse et al., 2021, 2016) and custom scripts developed based on Kegler et al. (2022). A TRF performs a linear transformation between one or more stimulus features and the corresponding EEG responses. This relationship can be mathematically represented as:

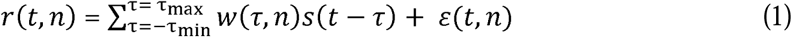

where the *r*(*t*,*n*) is the instantaneous EEG response to the stimulus at time t and channel n.

Here, *s*(*t*) is the absolute of the first derivative of the fundamental frequency |F0’|, and E(t) is the residual. The relationship between the response and stimuli is described at a certain range of time lags τ by the TRF weight w(r). The TRF weight, *w*(τ), can then be estimated by minimising the error between the recorded EEG responses and the predicted responses as below:

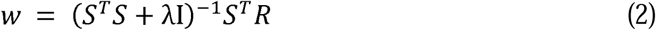

where S is the lagged time series of the stimulus. The. λ is the ridge parameter which is defined as.λ_n_ e_m_, where.λ_n_ is a normalized regularization parameter and e_m_ is the mean eigenvalue of the covariance matrix (Kegler et al., 2022). We used a fixed normalized regularization parameter of.λ_n_ = 0.1 for all participants. The time lag used to compute the relationship was-200 ms to 500 ms. This range was chosen based on a literature review (results shown in Appendix 1), where the peak latencies reached over 400 ms. The forward model was computed for all participants. The TRF weights were averaged across participants for the group analysis.

#### 2.4.5 Statistical analysis

Group-level statistical significance of the results was assessed with non-parametric permutation testing (1,000 permutations). For each permutation, the stimulus time series was time-shifted by a random value with respect to the EEG data, to abolish any meaningful time relationship between stimulus and response data, whilst preserving all other data features.

Any surplus stimulus data beyond one end of the EEG data (start or end) was moved to the other end. These misaligned stimuli were used to compute the TRF forward model for the permutation. Individual TRF weights of channel Cz were averaged across participants. The channel was chosen based on a previous study on single-channel analysis using AFG and SIN stimuli (Guo et al., 2022). Null distributions for TRF weights were created for individual datasets by taking the maximum absolute value across the TRF time series in each permutation (with the 50^th^-largest value of 1,000 permutations constituting the threshold for detecting significance at an alpha level of 0.05). The peaks of the TRF waveforms of the two AFG conditions as well as their performance were used to correlate with SIN performance using a bivariate Pearson correlation method.

After obtaining a model estimate for all datasets, the quality of the models was assessed by computing the EEG reconstruction accuracy, which is Pearson’s correlation between the predicted EEG output of the model and the real EEG data (Crosse et al., 2021). The reconstruction accuracy reflects how well the TRF models capture the encoding of the stimulus. This was compared to the permuted distribution (obtained the same way as described for TRF null distribution) with a pairwise t-test. A two-way (2×2) Analysis of Variance (ANOVA) was also performed on the reconstruction accuracy across two factors: stimulus type (SIN vs. AFG-F0 vs. AFG-1/F) and frequency bands (delta vs. theta). The correlation between the TRF peak amplitude of the AFG condition and SIN d’ was checked with Pearson correlation.

#### 2.4.6 TRF Source Localisation

Previous neuroimaging studies (Holmes et al., 2021; Teki et al., 2016, 2011) demonstrated high-level brain activities outside the primary auditory cortex at the superior temporal sulcus and the intraparietal sulcus for effects of duration and coherence of the figure. In the current study, tracking the frequency patterns during sound segregation could involve potentially different processing mechanisms distinct from pure figure detection in noise. Source localisation was used to explore if the locations driving the surface TRF activities were consistent with previous findings and if they were comparable to speech processing in noise.

This analysis was carried out using standardised low-resolution brain electromagnetic tomography using the using the MNI-152 template (sLORETA, version 20081104) (Pascual-Marqui, 2002). No individual structural MRI data were acquired for this study. The sLORETA provides a solution (5 mm spatial resolution of 6239 voxels) to the inverse problem at the cortical and hippocampal regions. The significant TRF peaks averaged across participants were transformed into MNI space and tested against the null distribution at the lags of the first and second peaks (Table 1). A one-sample t-test was used to compute p-values at each voxel, and the results were corrected with Bonferroni correction at the 0.05 alpha level.

**Table 1.** TRF peak time points chosen for the source localisation. These are the group-averaged latencies for the three conditions.

|  | Delta |  |  | Theta |  |  |
| --- | --- | --- | --- | --- | --- | --- |
|  | SIN | AFG-F0 | AFG-1/F | SIN | AFG-F0 | AFG-1/F |
| Peak Latency Early | 70 | 80 | 60 | 90 | 110 | 90 |
| Peak Latency Late | 260 | 280 | 220 | 180 | 210 | 170 |
| Peak Amplitude Early | -0.101 | 0.117 | 0.011 | -0.046 | 0.016 | 0.015 |
| Peak Amplitude Late | 0.116 | -0.081 | -0.005 | 0.047 | -0.015 | -0.012 |

## Results

### Performance on the active tasks

The d’ results are displayed in Figure 4. All three conditions achieved a good level of detection sensitivity (AFG-F0: mean (M) = 2.099, standard deviation (SD) = 0.979; AFG-1/F: M = 2.135, SD = 0.941; SIN: M = 1.971, SD = 0.441). No significant mean difference was found between all conditions.

**Figure 4.**
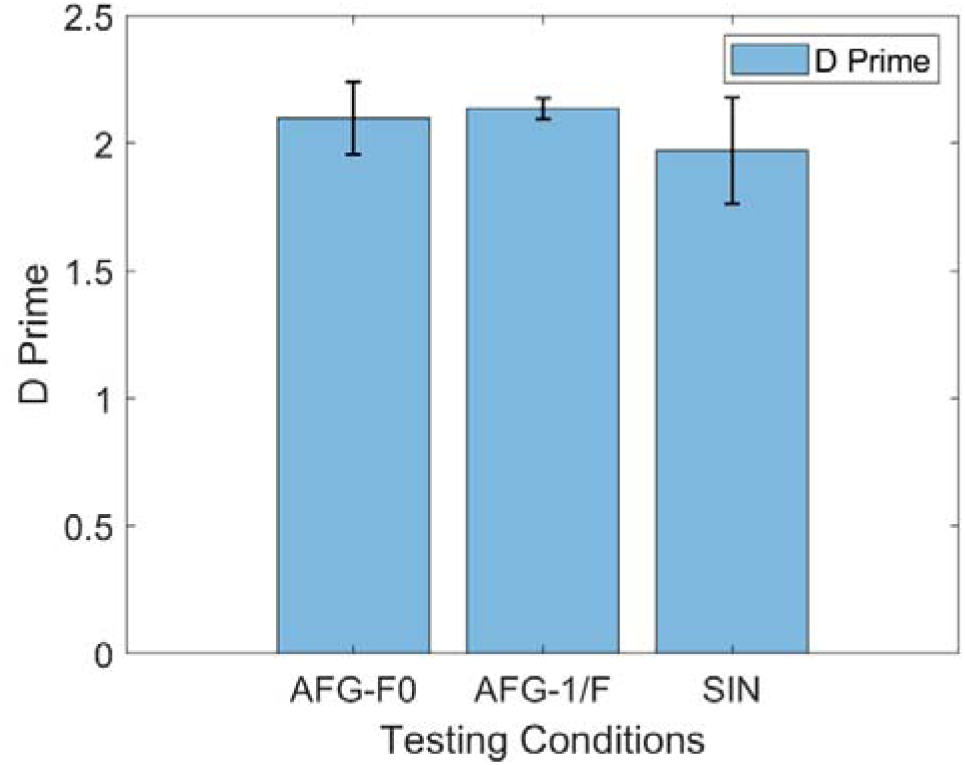
Participants’ performance in the experiments. The x-axis of the bar plot shows the three conditions as labelled, and the y-axis shows the d’ values. The black error bars show the standard error of the mean.

### Neural Responses at the Fundamental Frequency with a Single Channel

We examined neural entrainment to the contour of the fundamental frequency in the three experimental conditions by looking into the TRF weights obtained from the forward model on a group level. First, the responses for the SIN condition are shown in Figure 5a. The delta band analysis showed a significant early response from 20-110 ms which peaked at 70 ms. This was followed by a later positive response from 180-350 ms that peaked at 260 ms at Cz. The theta-band responses showed a narrower early response range from 80-110 ms that peaked at 90 ms. A significant late response from 160-190 ms that peaked at 190 ms was also observed. The scale of the TRF response was larger for the delta band than the theta band.

**Figure 5.**
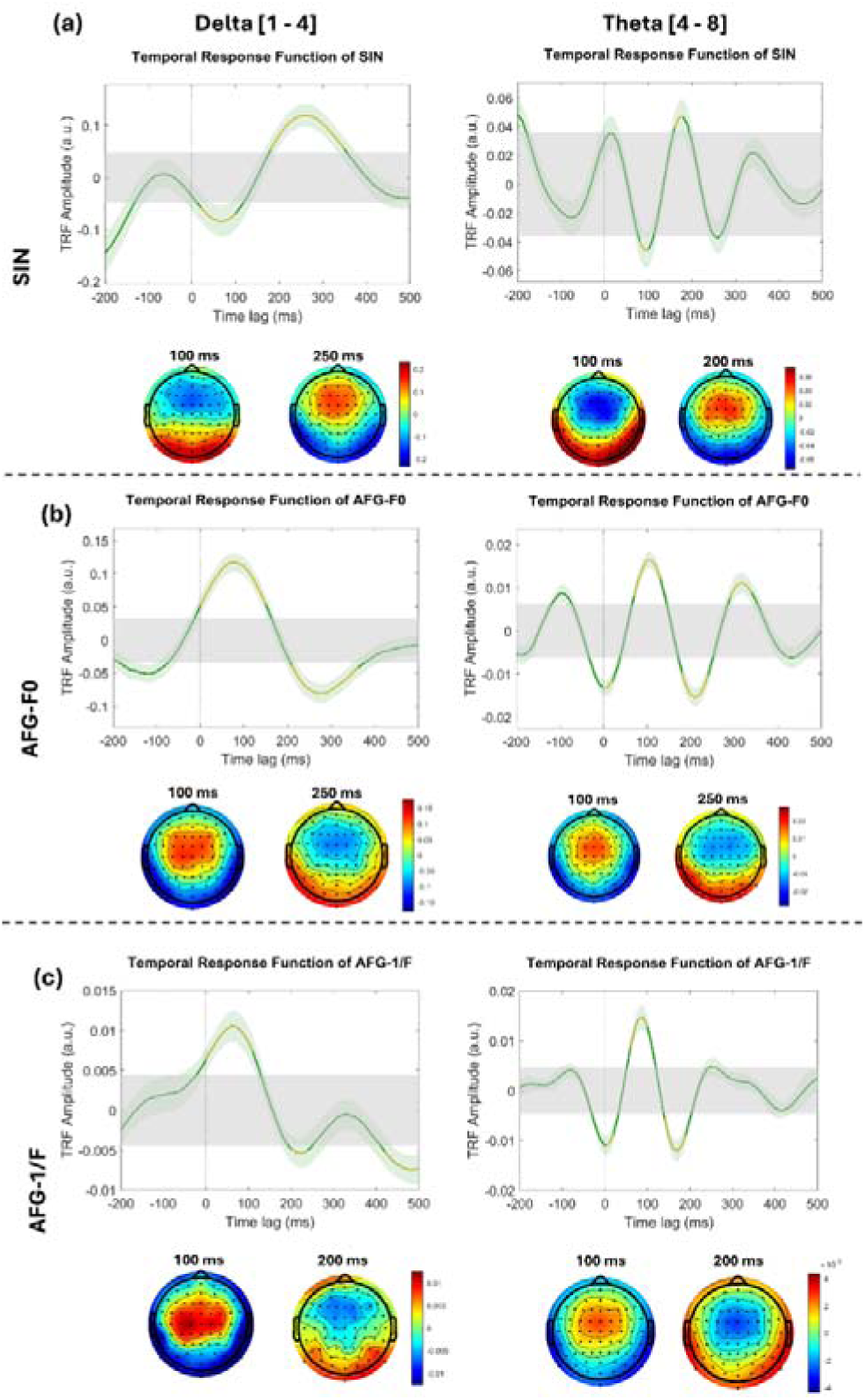
TRF responses and topographies for SIN (Figure 5 (a)) AFG-F0 (Figure 5(b)) and AFG-1/F (Figure 5(b)) at two frequency bands. The x-axes of the TRF plots show the time lag in milliseconds and the y-axes show the TRF weights in arbitrary units. The TRF waveforms are plotted in a dark green curve with a light green shadow as the standard error. The grey rectangular shadow marks the area of null distribution at 0.05 alpha level, and the yellow curves highlight the significant peaks. The left panel shows the delta-band TRF responses, and the right one shows the theta band.

The topographies of both frequency bands showed either negative or positive activities maximal at the frontal-central electrodes.

The TRF responses for the AFG-F0 condition are shown in Figure 5b. The delta range for AFG-F0 showed comparable magnitude with the SIN delta condition but the theta range was much smaller (the mean absolute amplitude of AFG-F0 in theta was more than three times smaller than that of SIN). The delta-band response showed a significant response before 150 ms that peaked at 80. The second peak was observed at 280 ms (range 210 ms – 360 ms). The early peak was also found with the theta condition with a range from 70 ms to 130 ms peaking at 110 ms, but the second peak happened earlier compared to the delta band, which was at 210 ms (ranged 180 ms – 240 ms). A third transient peak at 320 ms was also visible for the theta condition (range 290 ms – 340 ms).

The AFG-1/F condition (Figure 5c) showed a similar pattern to the AFG-F0 condition but with a much smaller magnitude in the Delta band. In addition to the first positive response before 110 ms peaking at 60 ms, and the second 220 ms peak (range: 20 ms – 24 ms), there was a later significant negativity at 420 – 540 ms in the delta band that peaked at 480 ms. Theta band showed significant 90 ms (range: 60 ms – 100 ms) and 170 ms peaks (range: 140 ms – 190 ms) similar to AFG-F0 and a transient third peak at 250 ms.

The TRF peak values of the early and late waves were extracted from the AFG conditions. We ran the Pearson correlation between the SIN d’ and AFG peak values of the Delta frequency (all data were normally distributed). We found a significant correlation between the early peak of AFG-F0 with SIN (r =-0.38, p = 0.037) but not the late peak (See Figure 6 for more details). The correlation between the AFG performance or TRF peaks of AFG-1/F and SIN was not significant (p>.206).

**Figure 6.**
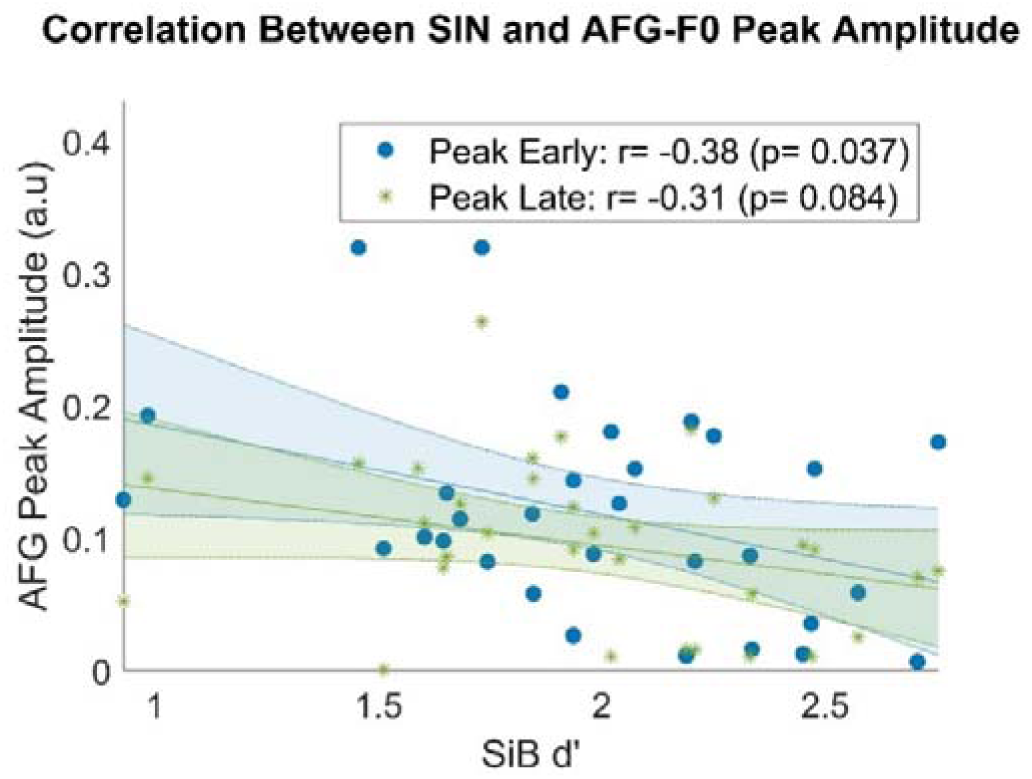
Scatterplot of the relationship between SIN d’ and the absolute TRF peak amplitudes of AFG-F0 at the early and late peaks (80 ms and 280 ms). The x-axis shows the d’ of SIN, the y-axis shows the peak values of TRF waveforms of the AFG-F0 condition. The shaded area plots the 95% confidence bounds. The legend shows the Pearson correlation coefficient (r) and its p-value.

The reconstruction accuracies of the TRF forward models are shown in Figure 7. All accuracies were significant compared to the null distribution (Table 1). The ANOVA results indicated a non-significant main effect of stimulus type (F (2, 62) = 1.90, p =0.158, effect size: ηp^2^ =.06). The main effect of frequency bands was significant (F (1, 31) = 139.94, p <.001, ηp^2^ =.82) due to the lower accuracy of the theta band (R_delta_ = 0.04, R_theta_ = 0.03). The interaction between stimulus type and frequency bands was significant (F (2, 62) = 17.23, p <.001, ηp^2^ =.36). The interaction was followed up by paired samples t-tests based on the descriptive data (Table 2) which showed lower predictive accuracy of the theta band in the SIN condition compared to AFG-F0 (t (31) =-5.72, p <.001), and AFG-F1 (t (31) =-4.83, p <.001), whereas there was no significant difference between the two AFG conditions.

**Figure 7.**
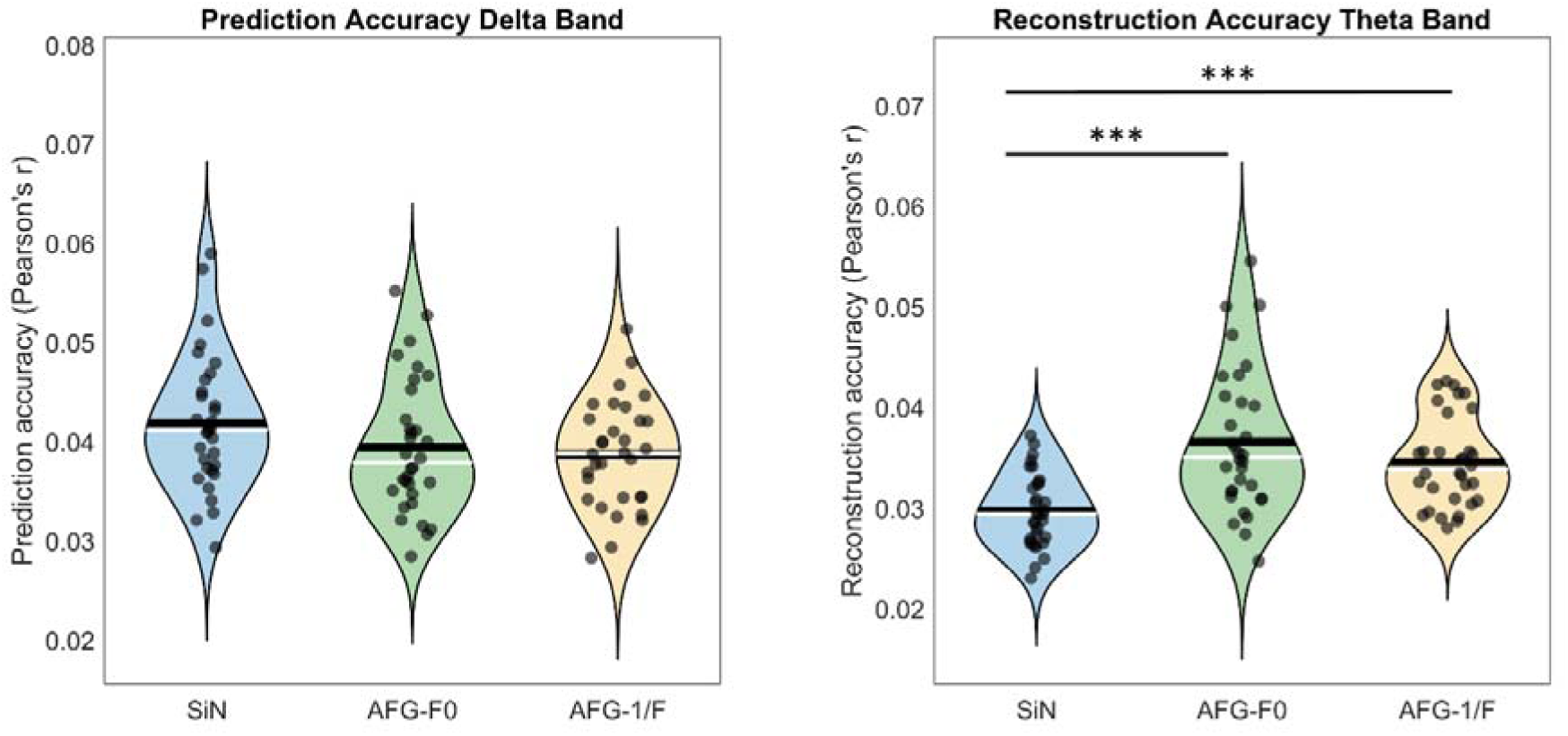
Prediction accuracies of TRF models in two frequency bands. The black dots show the reconstruction accuracy of individual participants. The median and the mean are plotted in white and black lines respectively. The asterisks with an underlying line illustrate the significant mean difference between conditions. Three asterisks ‘***’ suggest an alpha level of p<.001.

**Table 1.**
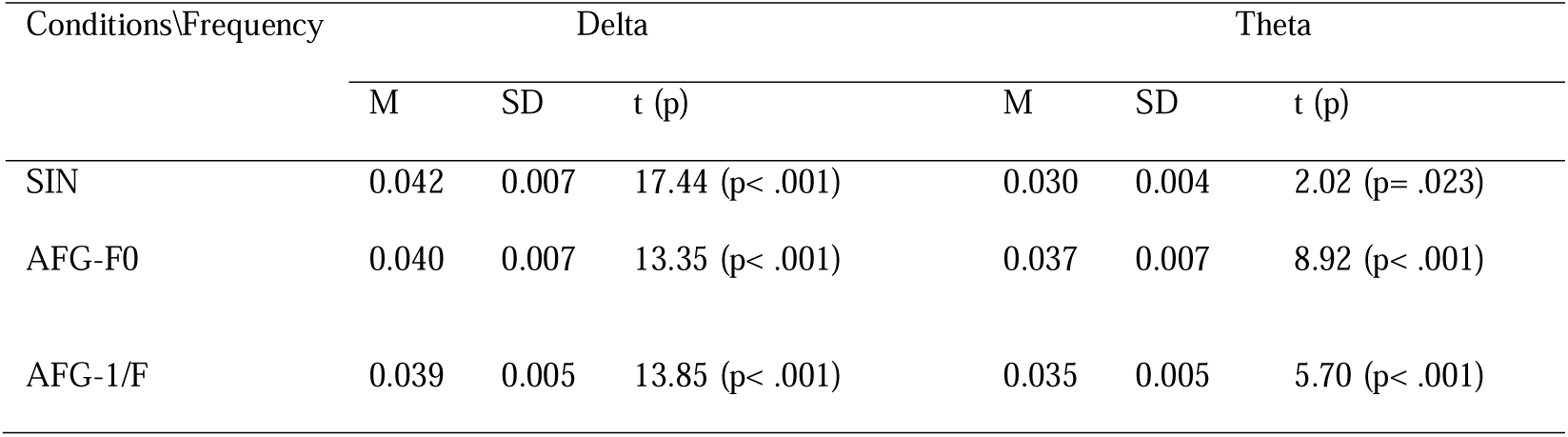
Prediction accuracies of TRF models in two frequency bands.

| Conditions\Frequency | Delta |  |  | Theta |  |  |
| --- | --- | --- | --- | --- | --- | --- |
|  | M | SD | t (p) | M | SD | t (p) |
| SIN | 0.042 | 0.007 | 17.44 (p< .001) | 0.030 | 0.004 | 2.02 (p= .023) |
| AFG-F0 | 0.040 | 0.007 | 13.35 (p< .001) | 0.037 | 0.007 | 8.92 (p< .001) |
| AFG-1/F | 0.039 | 0.005 | 13.85 (p< .001) | 0.035 | 0.005 | 5.70 (p< .001) |

In order to rule out the possibility of the lower SIN reconstruction accuracy being statistically introduced by our more stringent method of extracting the ‘pure’ pitch by regressing out the stress contour, we ran the analysis again with the stress contour left in the signal and found that while the accuracy did improve overall, the SIN condition was still significantly lower than the AFG conditions in the theta-band. This analysis can be found in the Appendix.

### Source Locations

As the reconstruction accuracy showed that theta-band tracking is significantly less accurate than the delta-band, EEG source analysis was only conducted on the delta condition. The sLORETA source analysis provided clear localised activities in the SIN and AFG-F0 condition but not the AFG-1/F. The peak time points are summarised in Table 2. The SIN and AFG-F0 TRF early delta peaks localised over the superior temporal gyrus (STG), middle temporal gyrus (MTG), and inferior temporal gyrus (ITG) (Figure 8). Bilateral source locations were seen over the medial temporal lobe (MTL: hippocampus and parahippocampal region) and insula as well. Outside the temporal lobe, activities over the prefrontal lobe, inferior frontal gyrus (IFG), medial frontal lobe, precentral gyrus and postcentral gyrus, superior parietal lobe (SPL), precuneus, and cuneus were found for both the AFG-1/F and SIN conditions. The SIN peaks showed a visually more lateralised pattern of activities compared to the AFG-F0 condition. The AFG-F0 first peak showed bilateral tracking but it became left-lateralised over the inferior temporal gyrus, parahippocampus, and the precentral gyrus.

**Figure 8.**
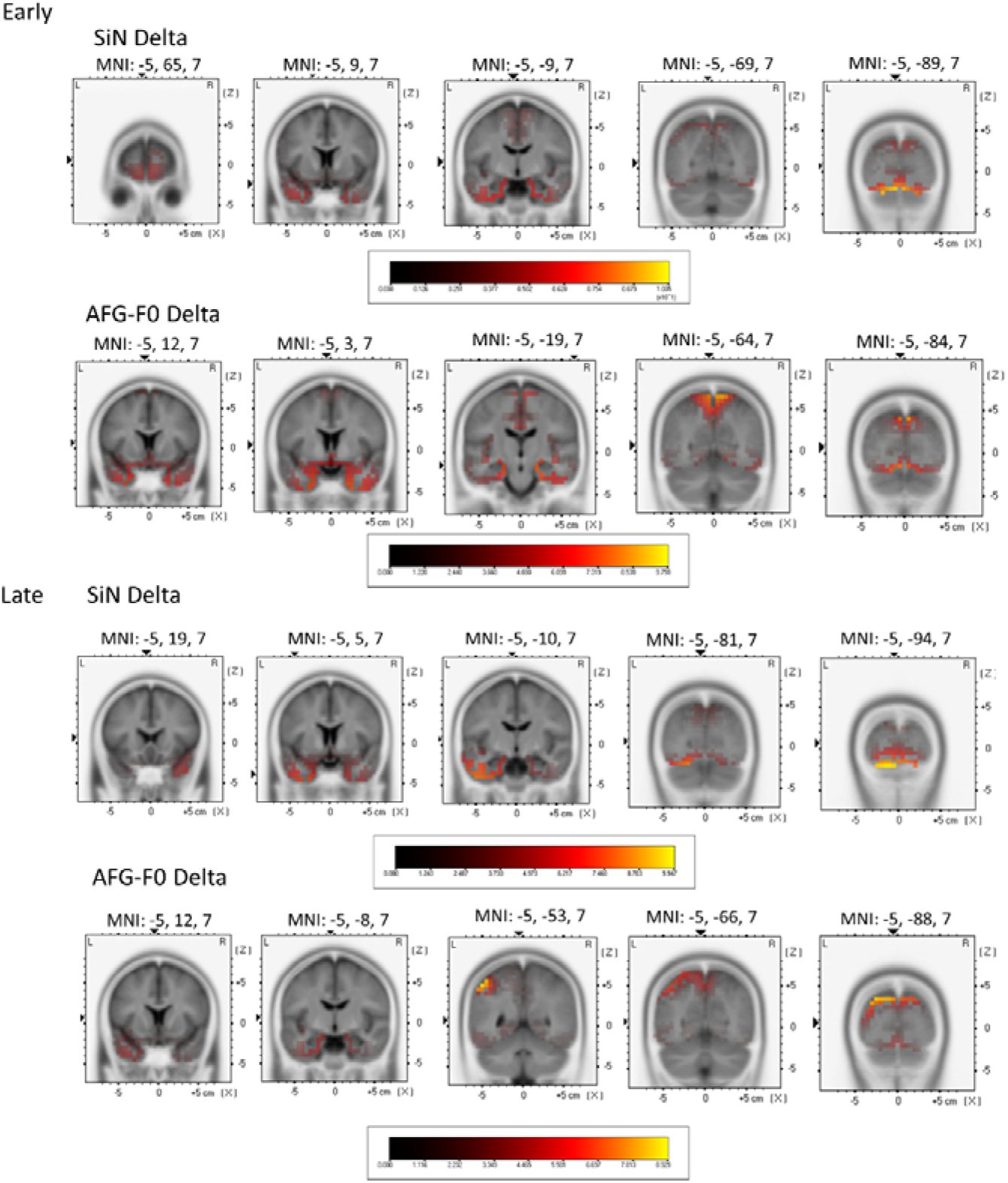
EEG source-level neural activities of Delta band TRF peaks to the fundamental frequencies in SiN condition and AFG-F0 condition.

## Discussion

The behavioural results showed that participants achieved excellent performance for all three tasks. As the design of the task required constant tracking of the target sound, the high performance indicated that the participants were maintaining their focus on the target sound. The TRF analysis showed that the brain can reliably track changes in F0 of human speech or F0-like frequency contour when masked by noise. However, different TRF morphologies and reconstruction accuracies were found both in terms of the type of stimulus and the specific frequency ranges that characterise neural oscillations in EEG.

### Similarity in cortical tracking and source locations of synthetic pitch-in-noise and natural SIN

The similarity between SIN and AFG processing has been demonstrated in both behavioural and neuroimaging domains previously with the stochastic figure-ground stimulus (Holmes et al., 2021; Holmes & Griffiths, 2019; Schneider et al., 2018; Teki et al., 2011, 2016).

O’Sullivan et al. (2015) further explored the neural tracking to the temporal coherence level of a random-frequency dynamic figure-ground stimulus and speculated that the pattern of TRF responses to AFG could be similar to that of SIN. Our results support this assertion, as detailed below.

Firstly, all testing conditions showed significant reconstruction accuracy based on the TRF forward model. This means that the brain can successfully entrain to the frequency change in either type of stimulus regardless of linguistic content, levels of predictability, or frequency range of neural oscillation. In terms of the TRF waveforms, the SIN and the AFG-F0 condition also demonstrated similar temporal encoding, with both conditions showing a peak at ∼100 ms and a second peak with inversed polarity at ∼250 ms in the delta band, and the same pattern at ∼100ms and ∼200ms in the theta band. These peak latencies were similar to previous studies that looked at TRF responses to SIN (Aljarboa et al., 2023; Bachmann et al., 2021; Ding & Simon, 2012). The similarity in the TRF time signature means that the brain likely is responding to the two stimuli on the same timescale, although the type of responses is not necessarily the same. The detailed TRF morphology will be discussed in the later section.

Further investigation into the source of the significant peaks showed that both SIN and AFG-F0 had generators over the temporal neocortex, parietal cortex, and medial temporal lobe, which were consistent with previous neuroimaging data (Holmes et al., 2021; Teki et al., 2016). In particular, parietal activities were found in both SIN and AFG conditions. Based on visual inspection, the left superior parietal lobe activities also seemed to be stronger and more widespread for figure-ground than speech-in-noise stimuli for the early peak, which was also supported by other studies comparing speech-in-noise to figure-ground or other non-speech signals in noise (Holmes et al., 2021; Kulasingham et al., 2021). The intraparietal sulcus (IPS), active during figure and speech tracking, has been implicated in stream segregation across multiple sensory domains due to its role in top-down attentional modulation (Calvert, 2001; Cusack, 2005). The engagement of medial temporal lobe shown here was also found in SIN before as well as another EEG study investigating the source of figure-ground segregation (Tóth et al., 2016). Studies have found that medial temporal lobe, particularly the hippocampus, is involved in not only auditory working memory but also extracting complex auditory patterns (See Billig et al., 2022 for a review on the role of hippocampus in auditory cognition).

### Polarity differences in TRF waveform of SIN and AFG

Unlike the wealth of literature on auditory-evoked responses (AEP), TRF research is relatively new, and the interpretation of the TRF forward model focuses mainly on comparing the absolute amplitudes or prediction accuracies between conditions, but the polarities are rarely discussed. However, the time lags and relative fluctuations in TRF waveforms can provide important information as well. The current study can offer insight into the interpretation of TRF morphologies. Firstly, we demonstrated that the neural tracking of SIN showed response patterns similar to the N1, P2/M200 responses (hereafter referred to as N1_TRF_ and P2_TRF_ to distinguish from the AEP components) in auditory-evoked potential, replicating previous findings (Aljarboa et al., 2023; Bachmann et al., 2021; Ding & Simon, 2012). The AFG condition, on the other hand, showed the opposite polarities at the same lags, which was also consistent with the previous findings on AFG stimuli (O’Sullivan et al., 2015a). The opposite polarities of the two conditions were reported by Horton et al. when they compared neural tracking of attended and unattended speech envelopes (Horton et al., 2013). They hypothesised that the inverted polarity seen in the unattended condition could reflect a suppression mechanism during auditory scene analysis, in which the attention network phase-locks to the inverse of the envelope of the noise. However, studies of a similar design did not find this pattern (O’Sullivan et al., 2015a; Power et al., 2012), although researchers did observe lower TRF amplitude and a degree of shifts in the latencies for the unattended stream. It is important to note that both the types of stimuli (speech vs. pure tone sequence, babble noise vs. tone cloud) used for the two conditions and the tasks (gap detection, repetition detection) were very different, which could result in this polarity inversion. It is therefore uncertain whether the polarity inversion we observed was related to attentional manipulation, and performance or whether it reflects the same neural process, but time-shifted.

However, literature on speech tracking in noise also suggests time shifting of TRF peaks as a possible explanation. We reviewed the recent literature on the neural tracking of continuous speech stimuli using TRF analysis or cross-correlation and found that a variety of peak latencies have been observed with very similar stimuli and filtering functions (see Appendix 1 for a summary of relevant studies).

The initial response to tracking a target speech can manifest as a positive TRF peak from 0ms to 160ms, or a negative peak from 0ms-200ms, followed by a peak of the opposite polarity of 100m-390ms (Behroozmand et al., 2015; Etard & Reichenbach, 2019; Brodbeck & Simon, 2022; Kegler et al., 2022; Muncke et al., 2022; Aljarboa et al., 2023; Panela et al., 2024). A few studies reported further fluctuations from 190 ms to 400 ms as well (Horton et al., 2013; Teoh et al., 2019). The majority of these studies reported results based on the amplitude of speech, but some of them used relative pitch similar to the current study (Bachmann et al., 2021; Brodbeck & Simon, 2022; Kegler et al., 2022; Teoh et al., 2019). This wide latency range of responses indicates that the definition of a TRF N1 P2 or M50, M100, M200 based on AEP could be misleading. Unlike the reliable N1 response in evoked potential, TRF literature does not necessarily show a 100 ms negative deflection. What literature shows is that TRF morphology can vary in the number of peaks and peak latencies with similar stimulus features used. The variation could be due to unknown task-specific effects, filtering function, or attention.

### Delta and Theta Band Encodes Different Levels of Acoustic Information

The ANOVA test on the model reconstruction accuracy found that neural tracking of frequency patterns on the theta band had significantly lower accuracy compared to the delta band. Furthermore, the post-hoc t-test showed that the lower SIN accuracy compared to the two AFG conditions also drove the interaction effect. The magnitude of the theta responses, however, exhibited the opposite pattern: speech tracking had a higher amplitude compared to AFG.

Cortical speech-tracking has been performed mainly on the speech envelope instead of pitch, as studies have found relatively low reconstruction accuracy for pitch encoding compared to acoustic envelope encoding, and even non-significant models for pitch in the theta band (Bachmann et al., 2021; Teoh et al., 2019). The current results, however, suggest that low pitch-encoding could be a speech-specific effect, as the AFG forward models held up their prediction accuracies. One possible explanation accounting for the relatively unreliable theta-band neural tracking for SIN is that the theta frequency might encode spectrotemporal information better than complex speech information. Previous literature has broadly related delta tracking to processing high-level features of speech, e.g. semantics and selective attention, whereas the theta band was linked to low-level acoustic processing such as rhythmic structure of speech (Ding & Simon, 2014; Zion Golumbic et al., 2012; Etard & Reichenbach, 2019; Peelle, 2013). The SIN condition used here encompasses high-level linguistic contents that can have an impact on the EEG responses whereas the AFG conditions only tap into sound segregation based on speech or speech-like pitch contours and harmonicity, which might be preferentially processed by the theta band with greater synchronisation between the neural signals and pitch information. On the other hand, the lack of linguistic information in AFG led to smaller TRF amplitude. This could be attributed to the effect of listening effort. Enhanced AEP N1 response has been observed for more effortful speech perception (Obleser & Kotz, 2011; Ghani et al., 2020).

### Relationship between EEG responses and behaviour, and potential clinical application

Finally, we found a significant negative correlation between the early peak of AFG-F0 and SIN performance but not the late peak. Traditionally, the correlation between the behavioural SIN and AFG performance has been demonstrated with large samples (n>100) (Guo et al., 2024; Holmes & Griffiths, 2019). We were therefore not expecting a significant correlation here between the performance of the two tasks themselves. However, we did find a significant negative association between AFG-F0 and SIN d’ with a moderate effect size, which could mean that the EEG neural tracking might be more sensitive than behavioural measures in showing this association. Higher amplitude for TRF weights can relate to a variety of auditory cognitive processes. A common finding in SIN perception is that attended streams tend to elicit stronger TRF responses, and the attentional modulation has been found to be strongest at ∼100–250 ms (Horton et al., 2013; Ding & Simon, 2012; Zion Golumbic et al., 2012). Higher demands for cognitive resources are posed for participants with lower SIN ability as they would need to recruit more attentional or working memory resources to compensate for their impaired fundamental sound grouping ability. A similar effect was found in speech processing in reverberant in a recent study, where the researchers combined pupillometry recording as well as EEG and found that listening effort and the strength of cortical tracking in the delta band increased with increasing difficulty in SIN perception (Ershaid et al., 2024). Enhanced AEP N1 response has been observed for more effortful speech perception (Obleser & Kotz, 2011; Ghani et al., 2020). While the current design cannot specify if the negative correlation shown here was due to general cognitive effect, future studies could incorporate measures of listening effort or attention to test the hypothesis. If the significant correlation can be replicated, the TRF signature of AFG can be potentially used to measure natural listening. The simple setup and efficient recording make it feasible for the development of its potential usage in clinics. This method has the advantage of posing intrinsically low demands on the patient’s ability to do complicated language tasks unlike most SIN tests in use at the moment and can dissociate the contribution of auditory processing and linguistic processing, which is influenced by language competence, education, accent, and other social factors.

No correlation was found between the peaks of AFG-1/F and SIN d’ despite the same level of reconstruction accuracy derived by the two AFG conditions. The difference could be driven by the distinct levels of stimulus-predictability. Natural sentence trajectories have a level of periodicity with regular recurrence of pitch patterns over time, but the 1/F pattern was mathematically generated and was not configured to have recuring similar pattern. The lower predictability led to sustained tracking on the target sound to facilitate figure-ground segregation for the AFG-1/F compared to other conditions as evidenced by the significant activities around 500 ms in delta. We also found pre-zero activities in the SIN and AFG-F0 conditions (not highlighted in the figure but fell outside the null-distribution, see Figure 5), whereas there was no significant pre-zero TRF peaks within the 200 ms window before zero for the AFG-1/F condition. The pre-zero activities for the natural-speech and AFG-F0 conditions were likely generated by correct prediction of upcoming pitch contour changes, which were not present for the AFG-1/F condition. This suggests that while an artificial pitch contour can generate the same level of model reconstruction accuracy, the underlying process might still differ from the processing natural speech contours.

To conclude, we have successfully demonstrated strong neural tracking of complex frequency patterns including natural pitch contour or speech-like contour in both SIN stimuli and AFG. The pattern of the pitch tracking showed similarity in the encoding accuracy, TRF peak latencies, and source locations in the Delta condition, but the AFG can obtain higher model accuracy than speech models in the theta band with lower magnitude possibly due to lower listening effort. The entrainment of pitch in the AFG stimuli also correlated with SIN performance, suggesting potential clinical use. A major limitation of this study is that the ‘performance’ measure of SIN processing was based on a simple task of detecting repetition with a small number of trials. This has possibly led to the small effect size of the correlation results, which would not be significant after applying correction for multiple comparisons. To obtain a more reliable relationship, a proper assessment of SIN performance is needed using speech-based tests and a large number of trials. Variability in age and hearing within the current sample may also influence the correlation results. While this diversity is important for ensuring a representative sample, a larger cohort is needed to validate the findings. Future experiments should be conducted to validate the correlation between the AFG-F0 amplitude and SIN performance. EEG recordings have limited spatial resolution. The generators of the figure-ground and speech tracking need to be validated with other methods with better spatial precision but similar temporal resolution such as electrocorticogram.

## Author contributions

Xiaoxuan Guo: conceptualisation, data curation, methodology, formal analysis, visualisation, writing - original draft, writing - review and editing

Guangting Mai: formal analysis, writing - review and editing

Yousef Mohammadi: formal analysis, writing - review and editing

Ester Benzaquén: conceptualisation, formal analysis, writing - review and editing

Kate Yukhnovich: writing - review and editing

Will Sedley (co-senior): conceptualisation, methodology, formal analysis, writing - review and editing, supervision

Timothy D Griffiths (co-senior): conceptualisation, methodology, writing - review and editing, funding acquisition, supervision

## Acknowledgements

This work was supported by the MRC [grant number MR/T032553/1].

